# Effects of mutations in pigeon *Mc1r* implicate an expanded plumage color patterning regulatory network

**DOI:** 10.1101/792945

**Authors:** Shreyas Krishnan, Richard L. Cryberg

**Affiliations:** Department of Biology, University of Texas at Arlington, Arlington, Texas, 76019

**Keywords:** *Mc1r*, pigeon genetics, pigmentation genetics, reverse genomics

## Abstract

Studies in mammals have shown that the Melanocortin 1 receptor occupies a pivotal role as a nexus for integrating paracrine and autocrine signals to regulate pigment production and type-switching between pheomelanin (red/yellow) and eumelanin (black/brown) pigment synthesis in melanocytes. Inactivating mutations in the *Mc1r* gene are responsible for recessive pheomelanic reddening traits in several species, while mutations that increase activity cause dominant eumelanic darkening traits in mammals and birds. Previous efforts to associate *Mc1r* coding variants with color variation in pigeons (*Columba livia*) have yielded conflicting results. Applying a reverse genomic approach, we discovered a novel 500 base pair frameshifting deletion in pigeon *Mc1r* that likely inactivates the single-exon gene. Segregation analysis revealed complete cosegregation (LOD = 12.2) with *smoky* (symbol *sy*), a recessive pigmentation trait reported in these pages by Willard F. Hollander 80 years ago. We coupled these findings with breeding tests to determine that *Dirty* (*V*), a dominant darkening trait, is allelic to *sy*, and identified two independent *V* alleles, one of which is associated with melanic morphs of two other bird species. In contrast to observations that *Mc1r* inactivation results in uniform pheomelanic pelage in mammals, its loss in otherwise wild-type pigeons occurs without apparent pheomelanism, instead increasing plumage eumelanism while leaving black bar pattern elements of the tail and wing largely intact. These findings require reconsideration of Mc1r’s presumed role in pigment type-switching in birds, and suggest the existence of Mc1r-independent pathways for eumelanic pigmentation pattern regulation unknown in mammals.

## Introduction

Coding sequence variation in the *Melanocortin 1 receptor* (*Mc1r*) gene has been found to be responsible for pigmentation traits in dozens of vertebrate species, primarily mammals and birds (Cieslak *et al.* 2011; Roulin and Ducrest 2013). The Mc1r protein is a seven transmembrane G protein-coupled receptor that occupies a key position in the melanogenesis regulatory pathway, directing melanin pigment synthesis through a cascade of downstream effectors via modulation of the second messenger cyclic AMP (cAMP) in response to extracellular ligands (Barsh 2006; Walker and Gunn 2010; Yang 2011). In mammals, the phenotypes of *Mc1r* variants may be described as falling along a spectrum of (eu)melanic dark morphs to pheomelanic red/yellow morphs. Mammalian melanic *Mc1r* alleles are dominant hypermorphs, typically single amino acid substitutions or short in-frame deletions involving the first extracellular loop or adjacent transmembrane spans that result in a constitutively active or hyperactive mutant protein, while pheomelanic alleles are recessive hypomorphs or complete loss-of-function (null) alleles; however, some null alleles result in near-white coats in certain contexts (Robbins *et al.* 1993; Klungland *et al.* 1995; Kijas *et al.* 1998; Våge *et al.* 1999; Schiöth *et al.* 1999; Newton *et al.* 2000; Everts *et al.* 2000; Ritland *et al.* 2001; Schmutz *et al.* 2002; Eizirik *et al.* 2003; Nachman *et al.* 2003; Fontanesi *et al.* 2010; Dürig *et al.* 2018). Either constitutively active or null *Mc1r* variants eliminate wild-type coat color patterns produced by eumelanin-pheomelanin pigment type-switching, typically resulting in solid black/brown or red/yellow coats, highlighting Mc1r’s pivotal role in the regulation of melanin pigment color patterns (Robbins *et al.* 1993; Fang *et al.* 2009; Kaelin and Barsh 2013). Among birds, color polymorphisms have been associated with over two dozen *Mc1r* variants in several wild and domestic species, but curiously, nearly all known avian *Mc1r* variants characterized thus far are darkeners; while a few variants that lighten portions of the wild-type plumage have been described, no solid red/yellow pheomelanic traits like those seen for null alleles in mammals are known to result from *Mc1r* variation in birds (Takeuchi *et al.* 1998; Theron *et al.* 2001; Kerje *et al.* 2003; Mundy *et al.* 2004; Doucet *et al.* 2004; Nadeau *et al.* 2006; Baião *et al.* 2007; Uy *et al.* 2009; Vidal *et al.* 2010a; b; Gangoso *et al.* 2011; Cibois *et al.* 2012; Johnson *et al.* 2012; Yu *et al.* 2013; Guernsey *et al.* 2013; Schmitt 2015; San-Jose *et al.* 2017; Janssen and Mundy 2017; Araguas *et al.* 2018; Kageyama *et al.* 2018).

Domestic pigeons harbor a vast diversity of pigmentary variation distributed among hundreds of breeds and varieties (Levi 1965). Although many genetic factors and interactions have yet to be rigorously dissected, classical genetic work by early pioneers, including Thomas H. Morgan and Willard F. Hollander, and continued efforts by hobbyists and amateur geneticists have resulted in the characterization of modes of inheritance and genetic interactions of dozens of Mendelian traits affecting plumage pigmentation and morphology (Morgan 1918; Hollander 1938, 1983; Sell 2012). These foundational contributions have enabled the application of modern genetic tools to propel advances in understanding processes underlying the unrivaled variation in melanin-based colors and patterns cultivated by fanciers over millennia (Levi 1963; Domyan *et al.* 2014). This kaleidoscope of pigeon color diversity also presents challenges due to the potential for abundant genetic heterogeneity and epistasis to wither associations for causative variants into a background of spurious signals generated by intense selection in small, structured sub-populations. Here we coupled reverse genomics with a family-based approach to avoid these pitfalls, mapping a novel frameshifting deletion in *Mc1r* to *smoky*, an autosomal recessive pigmentation trait with mild plumage darkening effects. We then leveraged this finding to establish allelism with *Dirty*, a dominant trait with grossly similar effects on plumage pigmentation, identifying two independent missense substitutions, Val85Met and Ser174Gly. Although seen in other studies of pigeon *Mc1r*, the effects of these widespread substitutions eluded previous researchers despite the former’s association with dark morphs in two other bird species (Mundy *et al.* 2004; Baião *et al.* 2007; Derelle *et al.* 2013; Guernsey *et al.* 2013). The lack of apparent pheomelanism and retention of major eumelanic pattern features in pigeons homozygous for the presumptive loss-of-function deletion allele call for a re-examination of assumptions regarding Mc1r’s role in avian pigmentation.

## Materials and Methods

### Animals and trait diagnosis

Pedigreed families were produced in our laboratory colony over a period of several years, with mated pairs housed in individual breeding pens. Feral pigeons were trapped in Tarrant County, Texas and humanely killed by CO_2_ asphyxiation prior to phenotypic assessment and tissue sampling. All animal procedures were conducted in accordance with protocols approved by the Institutional Animal Care and Use Committee of the University of Texas at Arlington (protocol numbers A09.009 and A14.004).

Phenotypic assessments were made blind to genotypes independently by at least two experienced evaluators under standardized light conditions. Evaluations were supported by observations recorded shortly after hatching, when the diagnostic value of certain phenotypic features (bill and skin pigmentation) are maximal. Individuals that could not be diagnosed due to epistatic piebald traits were excluded.

### Samples

Samples for genomic DNA isolation consisted either of whole blood obtained from live animals, or liver or skeletal muscle tissues collected from carcasses or late-stage embryos. For live animals, approximately 2 mL of blood was obtained by venipuncture and collected into EDTA vacuum collection vials. Unhatched fertile (determined by candling) eggs were collected 22 days after laying, and embryonic tissues were dissected for DNA extraction. Cells pelleted from whole blood or homogenized solid tissue samples were lysed by incubation with 20 µg/ml Proteinase K (reagents obtained from Sigma-Aldrich, St. Louis, Missouri, unless otherwise indicated) in 500 µL PK buffer (50 mM Tris-HCl, 20 mM EDTA, 100 mM NaCl, 1% SDS) at 55°C for 3–6 hours, and genomic DNA was extracted by standard phenol:chloroform:isoamyl alcohol extraction method (Sambrook and Russell 2006).

### Detection of variants from whole genome shotgun sequence libraries

Available sequence reads from whole genome shotgun sequence libraries from 40 pigeons of diverse breeds/varieties (described in Shapiro *et al.* 2013) were obtained from the Sequence Read Archive (SRA, BioProject accession PRJNA167554) and used to identify sequence variation within *Mc1r*. The SRA format was converted into split FASTQ files for mapping using the fastq-dump tool included in the SRAtoolkit provided by NCBI (https://github.com/ncbi/sra-tools/wiki/Downloads). These FASTQ formatted reads were mapped to the Cliv1.0 pigeon reference genome using the Novoalign short-read mapping software, version 2.08.02 (Novocraft Technologies, Selangor, Malaysia) with default parameters. Single nucleotide variants and short insertions and deletions were identified using the mpileup feature of SAMtools (Li *et al.* 2009). Simple sequence repeat length variations were characterized using the methods of Fondon et al. (2012), for repeats identified by Tandem Repeat Finder v2.30 (Benson 1999) with parameters “2 5 5 80 10 14 5.” Larger insertions, deletions, and other structural variation within 50 kilobases of *Mc1r* were identified by anomalous inferred insert size, mate-pair orientation, or mapped read depth retrieved using the mpileup feature of SAMtools (Li *et al.* 2009).

### Genotyping and DNA sequencing

Genotyping and sequencing primers were designed using Primer3 (Rozen and Skaletsky 2000). Deletion genotyping was performed using polymerase chain reaction (PCR): 1U *Taq* DNA polymerase (United States Biochemical, Cleveland, Ohio), 20 pmol of each primer (forward: CTTGACACTACGTGCTGTGG; reverse: ACATGCTGCGGAGGTGGT), 5 μL FailSafe 2× PCR premix G (EpiCentre, Madison, Wisconsin), approximately 25 ng genomic DNA template, and ultrapure deionized water to a final reaction volume of 10 μL. Genotyping thermal cycling conditions were an initial melt at 96°C for 3 min; followed by 30 cycles of 30 s each at 96°C, 61.7°C, and 72°C; with a 6 min final extension at 72°C. Sanger sequencing of the *Mc1r* coding sequence was performed in overlapping segments on PCR products (forward/reverse primer pairs GTGCCCTGGAGCTGAGGT / AGAGGAGGATGGCATTGTTG and GCGCTACCACAGCATCAT / CCATTATCGGTGTCCCACTG) amplified as above. Amplified products were treated with shrimp alkaline phosphatase and exonuclease I (both from United States Biochemical), and sequenced in both directions on an AB 3730xl capillary sequencer using BigDye 4.0 dye-terminator chemistry (Applied Biosystems, Foster City, California) by the UT Arlington Genomics Core. Electropherograms were visually inspected using BioEdit (Hall 1999).

### Statistical analysis

The significance of cosegregation and departures from Mendelian expectations were computed as exact binomial probabilities. The significance of departure of genotypic proportions from Hardy-Weinberg expectations in the feral pigeon sample was computed using Hardy-Weinberg exact test (Wigginton *et al.* 2005). Association of the *Mc1r* deletion allele with the smoky phenotype in feral pigeons was computed using Fisher’s exact test under a recessive model.

### Data availability

The authors state that all data necessary for confirming the conclusions presented in the article are represented fully within the article or are available in a public data repository. Whole genome shotgun sequence libraries are available from the NCBI Sequence Read Archive under BioProject accession PRJNA167554 (Shapiro *et al.* 2013).

## Results

Conducted as part of a genomic survey for variants of potential functional significance for pigmentary variation, an examination of available whole genome shotgun sequence (WGS) libraries for 40 domestic pigeons representing diverse breeds (Shapiro *et al.* 2013) revealed four *Mc1r* missense substitutions (Val85Met, Asp115Asn, Ser174Gly, and Arg217Cys), all of which have been reported previously (see Discussion). Unexpectedly, a stretch of approximately 500 bp with reduced mapped depth suggestive of a previously unknown intragenic deletion was observed within the *Mc1r* coding region for WGS libraries from several breeds (English carrier, English pouter, Egyptian swift, Indian fantail, laugher, Scandaroon, Spanish barb, and Birmingham, Oriental and parlor rollers), a surprising development given the multiple previous studies of pigeon *Mc1r* coding variation. To identify individuals possessing the deletion that could be used to experimentally confirm its molecular nature and inheritance, genomic DNAs from six individuals from our laboratory breeding colony selected based on the above breed distribution were screened by PCR using primers flanking the putative deletion. These screens revealed three individuals to be heterozygous for the deletion, including both members of a mated pair, and testing the offspring of the heterozygous parents confirmed normal Mendelian segregation (Figure 1A), supporting the conclusion that the deletion affects the endogenous *Mc1r* locus, and not a pseudogenized duplicate copy (Gross *et al.* 2017). Sequencing across the breakpoint confirmed a deletion of exactly 500 bp, corresponding to positions 5700250-5700749 of scaffold 123 (NW_004973333.1:g.5700250_5700749del), including a two-nucleotide site of microhomology indicative of a microhomology-mediated end-joining double-strand break repair process (Lieber *et al.* 2003). The deletion removes two bases of the fourth codon through codon 170, disrupting the reading frame and resulting in the predicted loss of all but the first three residues of the Mc1r protein (c.11_510del, p.Leu4HisfsTer100; hereafter *Mc1r^∆500fs^*; Figure 1B).

**Figure 1.**
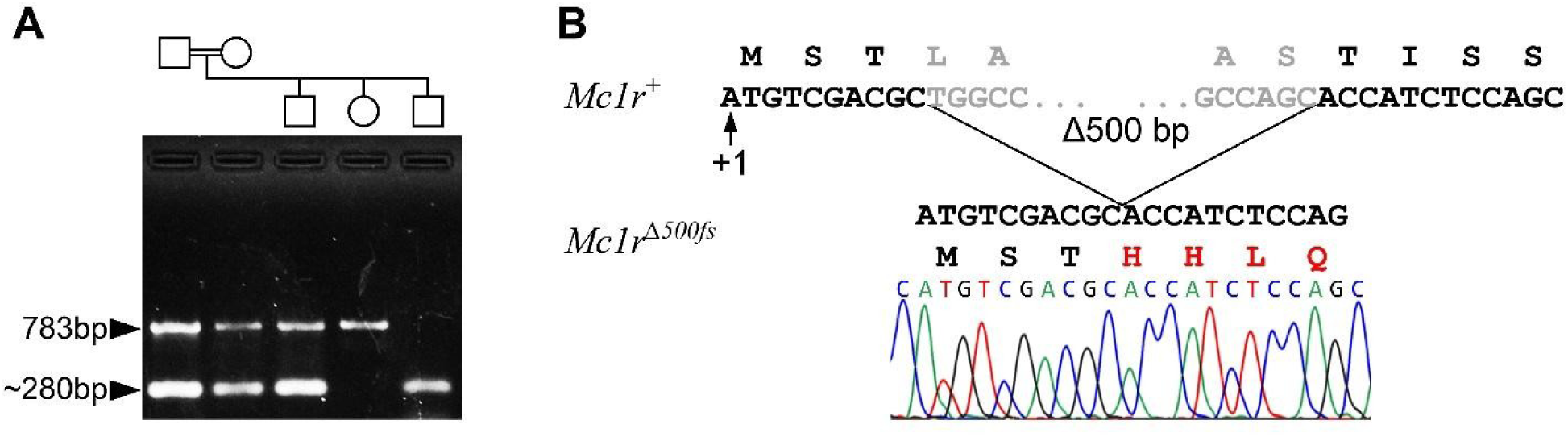
A. novel frameshifting deletion in *Mc1r* segregates in domestic pigeons. A. Screening by PCR identified apparent heterozygotes (left two lanes) for a putative ~0.5 kb deletion in pigeon *Mc1r*. The presence of all three genotypes among the offspring (right three lanes) of heterozygous parents demonstrates normal Mendelian segregation. B. Sanger sequencing of a homozygote revealed that the 500 bp deletion begins in the fourth codon, removes over half of the coding sequence, and disrupts the reading frame.

With the molecular nature of the deletion in pigeon *Mc1r* confirmed, we set about the reverse genetic process of attempting to identify any phenotypic effects. Activating mutations in *Mc1r* cause dominant melanic phenotypes in mammals and birds, while loss-of-function mutations produce recessive red/yellow coats in mammals, and are expected to have the same effects on plumage (Robbins *et al.* 1993; Klungland *et al.* 1995; Våge *et al.* 1997, 1999; Kerje *et al.* 2003; Ling *et al.* 2003). Pigeons possess one such trait, *recessive red* (symbol *e*), which would be an ideal candidate trait for the *Mc1r^∆500fs^* deletion had we not previously determined it to be caused by *cis*-regulatory mutations in *Sox10*, a downstream target of Mc1r signaling (Domyan *et al.* 2014). As there was no apparent pheomelanism in genotyped heterozygotes, other major pigeon reddening traits, including *kite*, *archangel bronze 1*, and *mahogany*, could be excluded on the basis of their modes of inheritance (Sell 2012). In the absence of obvious large-effect candidate traits, we adopted a family-based approach in order to limit the confounding impacts of epistatic factors and maximize sensitivity for traits of modest effect. Screening the founding pairs of pedigreed families from our laboratory colony identified two families segregating for the *Mc1r^∆500fs^* deletion, including a 14-member sibship from heterozygous parents (Family1, shown in Fig 1A), and a large three-generation filial pedigree derived from founders homozygous for alternate alleles (Family2). Both families were noted to segregate for *smoky*, a plumage darkening trait.

Characterized by W. F. Hollander in his doctoral dissertation research (1937, 1938), *smoky* (symbol *sy*; autosomal recessive) slightly darkens the light blue background of wild-type plumage, as well as the albescent (whitish) underwing and rump patch, creating an overall darkening impression very similar to that of the autosomal dominant *Dirty* (symbol *V*; Bol 1926) trait (Figure 2). Unlike *V*, which darkens essentially all pigmented regions, *sy* reduces the richness and definition of the dark wing bars, whitens the bases of pigmented feathers, slightly accentuates the blue tip edges of the rectrices distal to the black tail band, and reduces or eliminates the albescent strip on the lateral rectrices, causing these to become pigmented like other tail feathers (Bol 1926; Hollander 1937, 1938). Less noticeable in adults is the tendency of *sy* to reduce pigment intensity of the claws, beak, and skin, areas for which *V* has the opposite effect. The considerable variability in the expression of both of these traits and the abundance of epistatic interactions with other factors means that they are not unerringly recognizable across all genetic backgrounds; with experience, however, *smoky* can be reliably identified in many contexts, particularly when several individuals from the same stock are available for comparison (Hollander 1938; Sell 2012).

**Figure 2.**
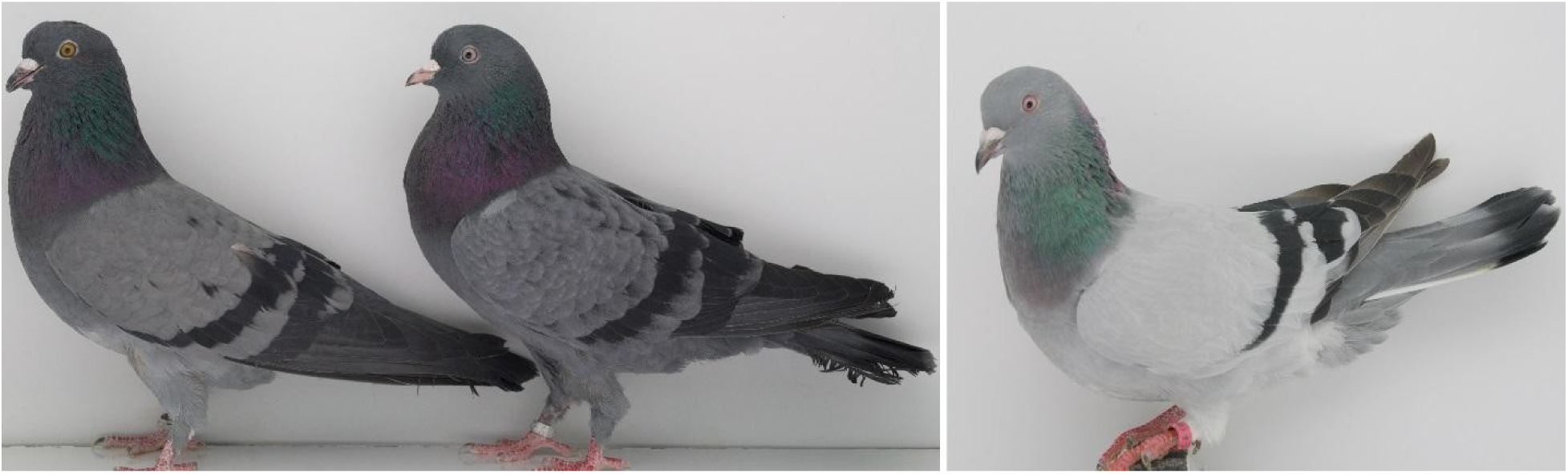
Typical effects of *Dirty* and *smoky* on plumage color are grossly similar. Both *Dirty* (*V*, left) and *smoky* (*sy*, center) darken plumage areas that are light blue-gray in wild-type (right), as well as the albescent underwing, producing overtly similar effects; however, unlike *V*, which darkens nearly the entire plumage, along with the skin, beak, and claws, *sy* lightens the integument, reduces definition of the dark wing shield pattern, and eliminates the whitish markings (albescent strip) on the lateral tail feathers (obscured by wing tip in the dirty individual). The increased width of the wing bars and black ticking on the wing shields of these *Dirty* and *smoky* birds relative to wild-type are due to variation at the *Checker* locus, which interacts additively with either *V* or *sy* (Hollander 1938).

In Family1, three smoky individuals were among 14 siblings obtained from a pair of non-*smoky* founder parents, identifying the founders as *sy* heterozygotes. In Family2, eight obligate carrier F_1_ offspring of a *smoky* male and non-*smoky* female P_0_ founders were mated *inter se* to produce 42 F_2_, including six *smoky* individuals. *Mc1r* genotypes of the founders of both families were consistent with *sy* locus genotypes, so we tested for cosegregation of the *Mc1r*^*∆500fs*^ deletion with *sy*. Independent phenotype assessments for *smoky* were made blind to genotypes for all 65 descendants of these founders by two experienced observers (S.K. & J.W.F.). Phenotype calls were fully concordant between observers, and genotypes exhibited complete cosegregation between *Mc1r^∆500fs^* and *sy*: all *smoky* individuals were homozygous for the deletion, while all other individuals had at least one copy of the full-length allele, and obligate carriers were heterozygous (Figure 3). The binomial probabilities of obtaining the observed complete cosegregation by chance, conditioned on founder genotypes, are p = 7.3 ×10^−5^ for Family1 and p = 7.8 ×10^−9^ for Family2, for a combined significance of p = 5.7 ×10^−13^ (LOD = 12.2).

**Figure 3.**
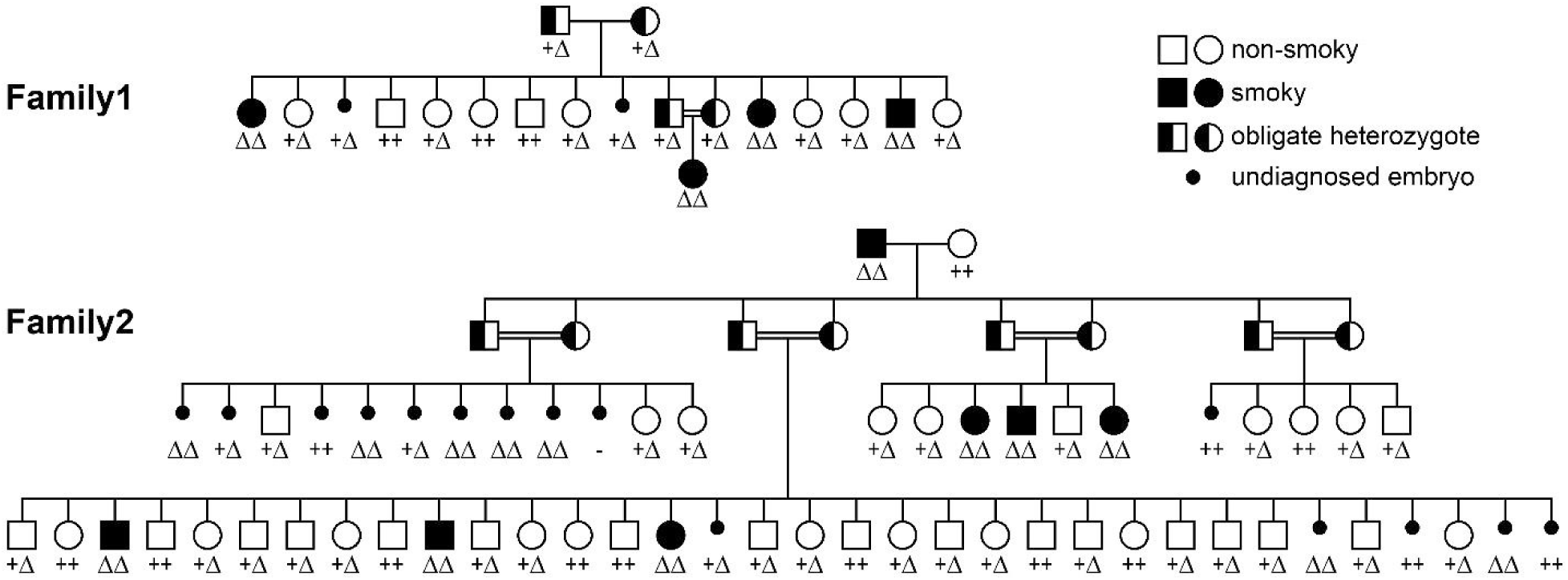
The *Mc1r^∆500fs^* allele cosegregates with *sy*. Segregation analysis in two unrelated families revealed complete cosegregation (p_Family1_ = 7.3 ×10^−5^; p_Family2_ = 7.8 ×10^−9^; p_combined_ = 5.7 ×10^−13^, LOD = 12.2). Embryos that failed to hatch are shown with their corresponding genotypes except for one that was unavailable (indicated by “-“). Unhatched embryos were not included in the cosegregation analysis.

Although Hollander’s original breeding data showed no evidence of reduced survival of *smoky* offspring (Hollander 1938), *smoky* appeared to be slightly under-represented in our mapping families. Only nine smoky offspring were obtained out of a total of 49 squabs hatched from heterozygote matings, but this under-representation was not significant (p = 0.18; binomial). While not included in the cosegregation analysis due to the uncertainty of phenotypic diagnosis of embryos, we also genotyped 16 F_2_ embryos that failed to hatch (Figure 3, small circles), seven of which were found to be homozygous for the deletion. The excess of *Mc1r^∆500fs^* homozygotes among unhatched embryos was also short of our chosen 0.05 significance threshold (p = 0.080), as was the excess of failures to hatch for deletion homozygotes in a combined analysis (p = 0.074). They were, however, substantial enough to motivate additional study.

A useful feature of pigeons as a model system is the availability of feral populations to provide insights regarding the fitness effects of variants of interest in natural populations of free-living animals (Johnston and Janiga 1995). Reduced early viability of *smoky* individuals would be expected to produce a deficiency of *sy* homozygotes relative to Hardy-Weinberg expectations (HWE) among feral adults. The high frequency of *sy* among domestic pigeons, the progenitors of feral populations, and Hollander’s observations of *smoky* individuals in feral flocks suggested that *sy* might be of sufficiently high frequency in feral populations that significant HWE departures could be detectable with smaller sample sizes (Hollander 1983). Screening 39 feral pigeons of unknown relationships for the *Mc1r^∆500fs^* deletion identified four deletion homozygotes, three heterozygotes, and 32 homozygotes for full-length alleles, a significant excess of homozygotes over HWE (p = 5.9 ×10^−4^; Hardy-Weinberg exact test). Although these data were not intended for genetic association testing purposes due to concerns about the reliability of trait diagnosis in pigeons of unknown ancestry, it is noteworthy that the four deletion homozygotes were the only *smoky* birds in the feral sample (p = 1.22 ×10^−5^; Fisher exact).

*Smoky* occurs in many pigeon breeds, and the representation of breeds possessing the *Mc1r^∆500fs^* deletion among the pigeon whole genome shotgun sequencing libraries is consistent with what is known about the breed distribution of *sy*. Breeds possessing the deletion include very old, morphologically disparate breeds with distant geographic origins, implying an ancient origin for the *sy* mutation (Levi 1963). A few breeds are reported to be fixed for *sy*, including carriers, barbs, magpies, and Egyptian swifts (Hollander 1938; Estabrook 2013). Three of these breeds were represented among the whole genome shotgun libraries, and constituted three of only six libraries that were homozygous for the *Mc1r^∆500fs^* allele: English carrier, Spanish barb, and Egyptian swift (p = 2.02 ×10^−3^; binomial).

Interestingly, the autosomal dominant *Dirty* (*V*) darkening trait was also segregating in both Family1 and Family2, exhibiting a peculiar inheritance pattern in which every individual expressed either *smoky* or *Dirty*. The probability of such an inheritance pattern resulting from random segregation of unlinked loci by chance is sufficiently small (p < 10^−12^) that it may be rejected, leaving two plausible explanations: either 1.) both families are fixed for *V* and epistasis of homozygous *sy* over *V* prevents the expression of *Dirty* in *smoky* birds (i.e. a two-locus model with epistasis); or 2.) *V* and *sy* are alternative alleles of the same locus (or tightly linked). Formally, the converse of model 1 (i.e. *V* is epistatic to *sy*, and it is *V* rather than *sy* that maps to *Mc1r*) could in theory also produce the observed inheritance pattern; however, the independent support for the association of *Mc1r^∆500fs^* with *sy* seen in the feral sample and the breed distribution data exclude this model. Although widely presumed to be separate loci by hobbyists, neither allelism nor epistasis between *sy* and *V* has been investigated, but their phenotypic similarities and the abundance of melanic variants in other species for which *Mc1r* has either been confirmed or excluded render either scenario plausible (MacDougall-Shackleton *et al.* 2003; Cheviron *et al.* 2006; Wlasiuk and Nachman 2007; Roulin and Ducrest 2013).

The availability of *smoky* F_2_ from Family2 offered a means to discriminate between the allelism and epistasis models using breeding tests. Under the two-locus model, all family members are homozygous for *V*, while under an allelism model *V* is excluded from the genetic constitution of *sy*/*sy* homozygotes. We performed testcrosses to distinguish between these scenarios. We mated two *smoky* F_2_ from Family2 to unrelated non-*Dirty* individuals that were (by happenstance) heterozygous for the *Mc1r^∆500fs^* deletion. Under the two-locus model, all offspring from such matings (*sy*/*sy*; *V*/*V* × *Sy^+^*/*sy*; *v^+^*/*v^+^*) will either be *smoky* or *Dirty*, but never wild-type. Alternatively, if the allelism model is correct, these *smoky* F_2_ cannot harbor *V*, and all offspring of these *sy*/*sy* × *Sy^+^*/*sy* matings will be either *smoky* or wild-type, but never *Dirty*. These test crosses produced a total of three *smoky* and six wild-type offspring, and no *Dirty*, establishing *sy* and *V* as alternative alleles of the same locus.

Melanic *Mc1r* variants identified in other species are most frequently single amino acid substitutions, or less commonly short in-frame deletions (Roulin and Ducrest 2013). In order to identify any coding sequence changes associated with *V*, we sequenced the entire *Mc1r* coding regions of the full-length alleles segregating in opposition to the deletion allele in both of the mapping families. Sequencing homozygous non-deletion F_1_ of Family1 showed that its founder parents were both heterozygous for the previously observed Gly174 variant (c.520A>G, p.Ser174Gly; *Mc1r^Ser174Gly^*) opposite the *Mc1r^∆500fs^* deletion. The *V*/*V* founder of Family2 was homozygous for the Met85 allele (c.253G>A, p.Val85Met; *Mc1r^Val85Met^*). Both of these variants occur in multiple pigeon WGS libraries (data not shown), and have been reported previously in other studies, as discussed below (Derelle *et al.* 2013; Shapiro *et al.* 2013; Guernsey *et al.* 2013; Ran *et al.* 2016).

## Discussion

While the array of color diversity amassed in myriad varieties of domestic pigeons offers great promise as a powerful resource for the genetic dissection of pigmentary regulation, the complexity of epistatic interactions and potential for genetic heterogeneity also present challenges (Domyan *et al.* 2014). We have combined reverse genomic and classical genetic approaches to isolate constituent Mendelian components of genetically complex pigmentary variation within families in order to avoid the pitfalls that have hampered previous efforts to link variation in pigeon *Mc1r* to pigmentation traits, uncovering insights that have the potential to alter our understanding of avian pigment pattern regulation. Our approach of working within well-characterized families served to bound the genetic complexity and enabled reliable genetic inference of similar and variably expressed traits, something not feasible for the diverse samples used for genetic association. After mapping the *Mc1r^∆500fs^* deletion mined from pigeon WGS sequences to *sy* by cosegregation, we leveraged the fortuitous presence of *V* in the mapping families to establish *V*-*sy* allelism, and identified two independent *V* alleles.

### Previous studies and breeder lore

The conclusions of the various studies that have sought to identify associations between *Mc1r* coding variants and pigeon color variation range from inconclusive (Ran et al. 2016), to excluding association (Miller and Shapiro 2011; Derelle *et al.* 2013), to marginally significant association of the Met85 allele with pheomelanism (Guernsey *et al.* 2013), to suggestive of association of the Gly174 allele with eumelanism (D. Smith, personal communication). The contradictions among the various studies are less substantial than some of their titles suggest, and we believe are largely attributable to methodological differences. The published studies either classified phenotypes by categorical appearance (e.g. pheomelanic vs. eumelanic), rather than on the basis of inferred genetic constitution for recognized traits, or considered only traits of large effect. The phenotypes of *Dirty* and *smoky* are sufficiently modest and background-dependent that under such schemes the classification of affected birds would be dominated by any large-effect traits with which they tended to co-occur. We suspect such a correlation is behind the curious findings by Guernsey et al. (2013) of a marginally significant association between pheomelanism and Met85, a substitution perfectly associated with the opposite effect (eumelanism) in red-footed boobies and lesser snow geese (Mundy *et al.* 2004; Baião *et al.* 2007). Breeders of red or bronze pigeon varieties routinely select for *V* and other darkeners as they are known to improve the appearance of red pigeons (Hollander 1937; Mangile 1973; Sell 2012). Modest non-random associations resulting from this selection are therefore not unexpected between *V* alleles and large-effect reddening traits, particularly in samples with substantial contributions from pigeons bred for exhibition. Because the robustness of their detected association (p = 0.03, uncorrected) contrasted so markedly with the much stronger associations typical of causative *Mc1r* variants, Guernsey et al. speculated that Met85 likely had only subtle or background-dependent effects that would require controlled crosses to resolve. Our results bear out these inferences, and our determination that Met85 is a major *V* allele brings new clarity to their results, returning pigeon pigmentation genetics to harmony with other birds (Mundy *et al.* 2004; Baião *et al.* 2007). The existence of the same substitution with similar effects in wild species validates pigeons’ utility as a model for understanding the genetic basis of variation in natural populations, and the surprising mildness of the *smoky* phenotype may help explain the paucity of *Mc1r* hypomorphs and null mutations identified in other birds.

The unpublished work of Dan Smith and colleagues at Goshen College differs from the other studies in that birds were scored for all known Mendelian factors, and the sample was restricted to a small number of breeds to limit potentially confounding genetic heterogeneity (D. Smith, personal communication). Such nuanced approaches to phenotypic classification offer clear advantages, but the complexity of pigeon pigmentary variation constrains their application in practice, as few traits are unerringly recognizable across most genetic contexts. Accordingly, epistatic traits precluded confident assessment of *Dirty* status for all but six individuals (4 *V*/–, 2 *v^+^*/*v^+^*), leaving their sample underpowered; however, the observed complete association of the Gly174 allele with *V* in these individuals, though short of significance, is supportive of our results (D. Smith, personal communication).

Breeders commonly credit the combined effects of *smoky* and *Dirty* for the quality of an individual solid-colored pigeon, a proposition seemingly incompatible with *sy* being both allelic and recessive to *V*. Such claims appear to be founded on conjecture divined from the traits’ separate effects in solid-colored backgrounds, rather than on rigorous genetic dissection; however, some may be more-or-less valid and yet still reconciled with our findings in either of two ways: First, breeders’ selection for intense coloration by design accumulates minor uncharacterized darkening traits, and may do so in a manner cumulatively indistinguishable from *V*, thus permitting concurrent homozygosity for *sy*. Second, like many nominally recessive traits, *sy* is not invariably fully recessive, and a tendency toward dominance becomes sufficiently pronounced in some solid-colored backgrounds that Hollander suggested breeding tests may be necessary to confidently discriminate *sy* homozygotes from heterozygotes in such backgrounds (Hollander 1938). Aesthetic superiority of *V*/s*y* heterozygotes over either homozygote could also explain the difficulties cultivators of some solid-colored varieties have encountered attempting to obtain intense, uniform coloration that will breed true (Sell 2012).

### Mc1r’s role as master regulator

The *Mc1r^∆500fs^* deletion excises over half of *Mc1r*’s 313 codons, including those encoding multiple transmembrane helices essential for G protein-coupled receptor stability and function (Rosenbaum *et al.* 2009). It also disrupts the reading frame, and with no potential site for termination-reinitiation in the remainder of the single-exon transcript, results in the predicted elimination of all but the first three residues of the Mc1r protein. We conclude that the *Mc1r^∆500fs^* allele results in the complete loss of Mc1r function.

Nonsense and frameshift loss-of-function alleles are abundantly represented among pheomelanic *Mc1r* variants in mammals (Robbins *et al.* 1993; Klungland *et al.* 1995; Newton *et al.* 2000; Beaumont *et al.* 2008; Fontanesi *et al.* 2009; Brockerville *et al.* 2013; Zhang *et al.* 2014; Dürig *et al.* 2018), but only a single presumptive null allele is found among the dozens of causative *Mc1r* variants reported in birds: Trp32Ter causes the melanic black-winged bronze (*b^1^*) trait of turkeys, which is also unusual among darkening *Mc1r* traits in that it, like *sy*, is recessive to wild-type (Vidal *et al.* 2010a). Another recessive avian *Mc1r* darkening variant is the Phe256del allele of melanic royal purple (*m*) guineafowl, the consequences of which for protein function are unknown, but could represent a partial, conditional, or complete loss of function (Vidal *et al.* 2010b). Interestingly, these three recessive darkeners each cause the elimination of small whitish secondary pattern elements occupying parts of individual feathers: the albescent lateral tail strip of smoky pigeons, the light wing bars of black-winged bronze turkeys, and the pearl spots of royal purple guineafowl. Although a requirement for Mc1r in the development of some apigmented regions may seem counterintuitive based on its G protein-mediated cAMP signaling activity, it is consistent with the finding that the whitish spots in developing feathers of guineafowl and some other birds contain unpigmented melanocytes maintained in an undifferentiated state by Agouti signaling protein (ASIP), an inverse-agonist of Mc1r (Lin *et al.* 2013). In addition to its well-known function to repress cAMP-stimulated eumelanin synthesis driven by Mc1r agonist binding, ASIP also inhibits murine melanocyte differentiation and proliferation via the Attractin/Mahogunin signaling pathway in a manner independent of cAMP second messenger signaling, but still dependent on high-affinity binding to Mc1r to support low-affinity interactions of ASIP with Mc1r-associated Attractin (He *et al.* 2001; Hida *et al.* 2009). We speculate that disruption of Mc1r-scaffolded cAMP-independent signaling by ASIP may be responsible for the loss of near-white secondary pattern elements in the smoky pigeon tail, black-winged bronze turkey wing, and royal purple guineafowl dorsum, and also note that the potential for some mutations to preferentially impact either G protein signaling or ASIP-Attractin scaffolding functions of Mc1r might account for some of the phenotypic variety of *Mc1r*-linked traits in birds, most notably gyrfalcons, barn owls, and chickens (Kerje *et al.* 2003; Johnson *et al.* 2012; San-Jose *et al.* 2017).

Since *Mc1r* null mutations eliminate dark eumelanic coat pigmentation in mammals, we were surprised to find the black bars of the pigeon wing and tail only slightly perturbed in *Mc1r^∆500fs^* homozygotes, indicating that the signals governing their expression are transduced largely independently of Mc1r. This apparent divergence from mammalian pigment pattern regulation contrasts markedly with the similarity of the phenotypic effects in pigeons of mutations in downstream melanogenesis genes responsible for the four major types of human oculocutaneous albinism (see companion paper), indicating that melanogenesis pathway endpoints are not radically different between the taxa. The resilience of some, but not all melanin-based pattern features to the *Mc1r* alleles reported here suggests that “pathway” is no longer an appropriate term for avian pigmentary regulation, and “network” is a more apt descriptor. The identity of one such parallel regulatory network node is suggested by the recent identification of *NDP* as the pigeon *c* locus, responsible for *barless* and *Checker* traits affecting only the wing bars (Vickrey *et al.* 2018). It should not come as a surprise that a more elaborate melanogenesis regulatory apparatus might be employed to decorate feathers than hairs, given the greater potential offered by a two-dimensional canvas (Mundy 2005; Lin *et al.* 2013); however, conserved roles for parallel regulatory nodes in mammalian coat patterning may yet exist.

### Mc1r’s role in pigment type-switching

Plumage color patterns formed by changes in pheomelanin-eumelanin proportions are common in birds and thought to be regulated by Mc1r and its ligands, as is the case for mammalian pelage (Yoshihara *et al.* 2012; Lin *et al.* 2013; Gluckman and Mundy 2017). Yet while some alleles may enhance pheomelanism in red genetic backgrounds, such as *recessive red* pigeons or wild-type red jungle fowl, no avian *Mc1r* variant identified thus far manifests the ability to independently affect wholesale switching of wild-type eumelanic plumage to the uniform red/yellow pigmentation typical of *Mc1r* null alleles in mammals (Mundy 2005; Boswell and Takeuchi 2005). That the iconic alternating pheomelanin-eumelanin bands of turkey tails are retained in black-winged bronze turkeys should serve as a high-visibility warning of something amiss with the conventional wisdom; however, the significance of this important finding has gone largely unnoticed, possibly discounted due to its uniqueness or the potential for post-transcriptional rescue (Vidal *et al.* 2010a). The identification here of a clearly inactivating, yet phenotypically mild, darkening disruption of *Mc1r* in pigeons affirms that the similarly mild, darkening *Mc1r* nonsense variant in black-winged bronze turkeys isn’t a red herring, and the possibility must be considered that the biological context of Mc1r in plumage pigmentation may deny it the capacity to independently spawn the types of uniform pheomelanic effects of the classical *extension* (*e*) locus *Mc1r* mutants of mammals (Kerje *et al.* 2003; Mundy 2005; Vidal *et al.* 2010a). The pigeon *recessive red* trait was symbolized *e* because it was expected to be homologous to mammalian *e* locus mutants (Metzelaar 1926), but mapping two pigeon *e* mutations to *cis*-regulatory element deletions in *Sox10*, a target of Mc1r signaling activity, demonstrated that this pheomelanizing capacity resides downstream of Mc1r in pigeons (Domyan *et al.* 2014). Perhaps Mc1r is not the fulcrum upon which pheomelanin-eumelanin switching hinges for feathers to the extent that it is for hairs. Far from a radical proposition, such a difference might entail little more than incremental shifts in baseline activity of downstream feather melanogenesis components to favor the synthesis of eumelanin over pheomelanin at any level of cAMP stimulation, rather than only at elevated levels as in mammals (Barsh 2006).

### Genetic symbols

Our findings call for some genetic symbol revision. In keeping with conventions for pigeons, we retain Bol’s earlier *V* and suggest *v^sy^* for the *smoky* allele of the *Dirty* locus in recognition of its newfound allelism to *V* and Bol’s priority (Bol 1926; Hollander 1937, 1938). The different genetic backgrounds of the families in which the *Dirty* alleles examined here segregated precluded meaningful assessment of potential differences in their phenotypic effects, so we will distinguish their symbols only by superscripted numerals. Since offspring from Family1 were unavailable for breeding tests, the association of the Gly174 variant with *V* must be considered somewhat tentative, as without such tests we cannot exclude the possibility that the melanism of non-smoky individuals in this family was influenced by background factors hypostatic to *smoky*. We propose symbolizing the Gly174 variant as *V^1^* in recognition of its prior preliminary association with *Dirty* by Smith and colleagues, and the Met85 variant as *V^2^*.

Our results should prompt reexamination of additional *Mc1r* variants from previous surveys in pigeons and other birds for associations with relatively mild phenotypes that may have been overlooked. More importantly, these findings offer new insights into plumage pigmentation regulation, and identify powerful new tools for the genetic dissection of pigeon pigmentary variation that can be leveraged to identify and characterize additional components of the avian melanogenesis regulatory apparatus.

## Acknowledgments

Thanks to John Fondon, and thanks to lab members for assistance with animal care and sample collection. This work was supported by institutional funding from UT Arlington (to J.W.F.).

Numerous discussions about pigeon trait diagnosis with expert breeders and genetics enthusiasts, including Kerry Hendricks, Joel Kinkade, Amy Klopman, Nicholas Fontenot, Michael Bordelon, and David Rinehart, were invaluable in guiding this research. We would also like to Dan Smith at Goshen College for sharing his unpublished results with us, and for helpful discussions.

